# Information Network Flux interprets the importance of regulatory circuits in the protein and gene spaces

**DOI:** 10.64898/2025.12.03.692218

**Authors:** Boxin Xue, Ran Jiang, Yue Xue, Hong Zhang, Sirui Liu, Yi Qin Gao

## Abstract

Cells receive information from environment, make decisions, and execute biological functions. This sensor-actuator controlling mode is common, and has benefits for system stability especially when there exist feedback-loops for action correction. In cells, transcription factors (TFs) play the role of “brain” since they control the expression levels of about 20,000 protein-coding genes. However, the detailed topological structure of the regulating relationships among them is far from clear. Circuit-like pathways, which usually contain protein-protein interactions (PPIs), TF-gene regulations and gene expressions, are pervasive in cells, however, their importance is yet to be systematically elucidated by unifying information at both protein and gene levels. Here we developed Information Network Flux (INF), an algorithm that could simulate information transmit on multi-layer networks, and integrate protein and gene level information for cell type specific pathways that respond to given perturbation. We used topological analysis to identify regulatory circuits and found that the TFs participating in these circuits, especially those acted as “sensor” or “actuator” are highly relevant to cellular response to external signals. At the gene level, circuit genes serve as efficient cell type indicators, which showed better performance than commonly used gene expression levels. At the circuit level, shorter circuits enrich housekeeping functions and longer circuits exhibit increased tissue-specificity. These findings suggest that these circuit pathways may contribute to the identity and stability of cell states.

## [Introduction]

The virtual cell model aims to replace expensive and complex single-cell experiments with digital simulation operations, and to explore the response of biological cells to various external stimuli in a high-throughput manner, thereby greatly speeding up the process of new drug development^1–4^. One crucial task for virtual cell modelling is to interpret in detail how cells respond to stimuli. With the rapid advancement of deep learning methods, several foundation models in different aspects have been developed^5–9^, together with models for special tasks^10–13^. Although a large number of AI models are being developed^14–18^, there are three remaining challenges. One is the lack of suitable experiment data (especially multi-omics paired data and time-series evolutionary data), which greatly limits the application to many special cases due to the lack of accuracy^19^. Another is the black-box nature of deep-learning-based modes, which limits the power of explanation especially when the signaling details are important. A third problem lies in the hierarchical dynamic nature of data which are thus of different importance in determining the cellular states, adding difficulties for pure data-driven approaches to decipher the cellular mechanisms. At the level of gene regulation, information primarily flows from transcription factors (TFs) to the downstream genes they regulate, resulting in a network with strong unidirectionality and aggregation. At the protein level, however, information can be transmitted bidirectionally through protein-protein interactions (PPIs), allowing it to propagate backward from functional proteins to transcription factor proteins, thereby forming information "circuits". Thus, protein-protein interaction networks and gene regulatory networks exhibit fundamentally distinct organizational architectures, which necessitates that in molecular mechanism analysis, we must not directly conflate a protein with the gene responsible for its expression. Thus, a “physical” model, which can explain detailed information in hierarchical dynamic nature for custom cases, is still needed. We speculate a functioning virtual cell model to be a “mixed” one containing three components: (1) AI agents that integrate emerging knowledge into a coherent and comprehensive framework; (2) A colossal of data-based AI models for special tasks as tools and (3) Explainable models from a molecular interaction network point-of-view. In this work we focused on the last component mentioned above. The crucial task here is to predict key affecting factors (e.g. important molecular nodes or interaction network structure). A cell needs to collect information, make decisions, and then take responses, in a sense, similar to a robot. In robot control, the sensor-actuator controlling mode is common, where feedback loops are formed for action correction. In this study, we sought to examine the existence of similar control patterns in cells. For a cell, it is commonly accepted that the TFs together formed its “brain” ^20–22^. We further classify functions of TFs into “sensors” and “actuators”. The “sensor” TFs collect information from the cell (and external) environment, for example, through the interaction between TF proteins and functional proteins to form TF protein clusters (TFpcs) which participate in different regulation processes. On the other hand, “actuators” receive information from “sensors” and send task to the functional proteins, e.g., through the expression regulation of target genes. If there exist, say, a circuit containing both a sensor and an actuator, then the actions from the actuator could be viewed as supervised by the sensor. Since circuits normally play important roles in network stability and are expected to be important for the identity and robust response of cells, we try to address the following questions through the current study: Does this kind of regulation pathway exist in real cells? What are the specific biological functions of TFs that act as sensors or actuators? What is the role of these circuits in the cell fate determination?

The Gene Ontology (GO) ^23,24^ is a standardized classification system that annotates gene products across species, organizing their functions into three interconnected ontologies: Molecular Function (molecular activities), Cellular Component (subcellular locations), and Biological Process (macroscopic pathways). By assigning hierarchical, controlled terms to genes, GO enables systematic comparison of genetic roles, facilitates data integration across platforms, and supports functional enrichment analysis to decipher biological mechanisms, disease pathways, and evolutionary relationships. Making use of this rich information, we develop here Information Network Flux (INF), an algorithm that can simulate multi-layer networks for cell-type-specific pathways that respond to given perturbation. We construct a multi-layer network containing both knowledge-based regulation information including PPIs and RNA-seq data, as well as the GO features for the Biological Process. We made an especial effort for INF to be explainable. Particularly, we used topological analysis to find circuit patterns. We identified “strongly” linked sensor-actuator controlling circuits in different cell types, which show explicit cell-specificity due to the difference in gene expression patterns. We applied GO analysis to TFs acting as sensors or actuators and found their primary roles to be responses to diverse stimuli and regulation of cell development. We found that these “circuit genes” are efficient cell type indicators, which perform better in classifying cell types than commonly used gene expression analysis. This comparison supports the important roles of regulation circuits and their associated genes in the cell fate determination. Finally, we proposed hypothetical models for explaining the correlation between the circuit and tissue-specific RNA expression as well as the circuit-mediated signal response.

## [Results]

### Information Network Flux (INF) on multi-layer cellular network

For a given cell state, we tried to investigate the perturbation caused by a given stimulus, which is a fast response process before cell state transformation. Since limited experimental methods can be applied to measure such fast responses without changing the cell state but exist a vast range of related knowledge, it is worthwhile to construct a theoretical model based on integrated knowledge. In this approach, custom PPI and TF-regulation networks were built combining both RNA-seq results and databases (Methods). We have developed a message-passing-based sampling method on the aforementioned network and named it Information Network Flux (INF). INF has two main components: (1) Initiating from each potential perturbation source, it simulates the information propagation process to predict the list of affected TFpcs and genes, forming a reduced regulatory network that exclusively encompasses TFpcs and genes, accompanied by a quantitative scoring system that describes the magnitude of impact. (2) By performing a topological analysis of the network obtained in (1), a list of circuits along with their corresponding sensors and actuators is derived, and the "circuit flow efficiency" of these circuits is scored.

For the first component above, information propagation starts from a given protein node as the perturbation source. The information subsequently propagates within the PPI network, with propagation values determined based on the knowledge-based PPI network skeleton^25^ and the corresponding RNA expression levels of each protein node^26,27^ (Methods, Figure 1. Ideally, protein levels are preferred for calculating the weights, but due to the lack of data, in the current model, RNA levels are used instead.) In most cases, the TF proteins bind to DNA with the help of other proteins in the form of protein clusters^28,29^. Therefore, we used both gene expression levels and GO when simulating the formation of specific functional TFpcs, which contained information with specific GO features. We categorize TFpcs based on all possible GO terms associated with the functional TF proteins they contain. Each category is defined as a “single GO” TF cluster, characterized by a specific GO term. The TF proteins responsible for executing this function are subsequently designated as ’core’ TF proteins. (Methods, Figure 1). Information is then transmitted to target genes through the “single GO” TFpc, weighted by regulation weights in the custom TF regulation knowledge graph. To simulate selective binding of TFpcs, only the target genes containing the same or successor GO terms in the GO tree (Methods) are considered in the calculation of the total sum weight. After sampling from all 18,710 proteins (those with extremely low expression were omitted), we obtained a message-passing network containing only genes and TFpcs. These networks contain cell-type-specific signaling pathways, which are enriched with information content and are thus easily explainable. Within the multi-layered gene-protein interaction network, the aforementioned TFpcs serve as an “intermediate layer” to connect between the gene layer and the protein layer. Initiating from any gene, the signal propagates from the gene layer to the protein layer through the successful expression of the corresponding protein, subsequently transmitting through PPI networks to TFpcs. These TFpcs then regulate the expression of downstream genes within the gene layer, ultimately forming a closed information loop. For the second component, by analyzing the interaction network encompassing TFpcs and all genes, we wish to enable the precise definition of ’circuits’ as well as ’sensors’ and ’actuators’ within the system. We first considered DiG_signet, a reduced directed graph with only genes and “single GO” TFpcs (Figure 2a). The edges in DiG_signet are of three types: (1) TF regulation from “single GO” TFpcs to target genes (2) Gene expression regulatory relationships from TF genes to the TFpcs where the expressed TF proteins serve as core proteins (3) Influence from all genes to “single GO” TFpcs (obtained from sampling of highly expressed proteins as stimulus, named as “GPCI signaling” for short in the following). Based on this interaction network, we defined “sensors” and “actuators”: (1) The TF proteins, which formed “Single GO” TFpcs by hierarchical binding of other proteins, “sense” information from the cytoplasm environment, thus are regarded as sensor candidates. (2) All TF genes can be regarded as actuator candidates, since they express TF proteins to regulate target genes or modify the function of other TFpcs through PPI. (3) TF genes involved in circuits containing sensor candidates are further classified into sensors and actuators respectively. Under this definition, sensors collect information and then transmit it through a series of regulatory interactions to effectors. These effectors relay signals to functional proteins to carry out their functions, while feedback signals are transmitted back to the sensors via GPCI signaling. We sampled all sensors and actuators through finding all possible circuits between all sensors and actuators candidate pairs (Methods). For each cell type, each circuit’s "circuit flow efficiency" score as well as the dominant paths between them were defined and calculated (Methods).

**Figure 1.**
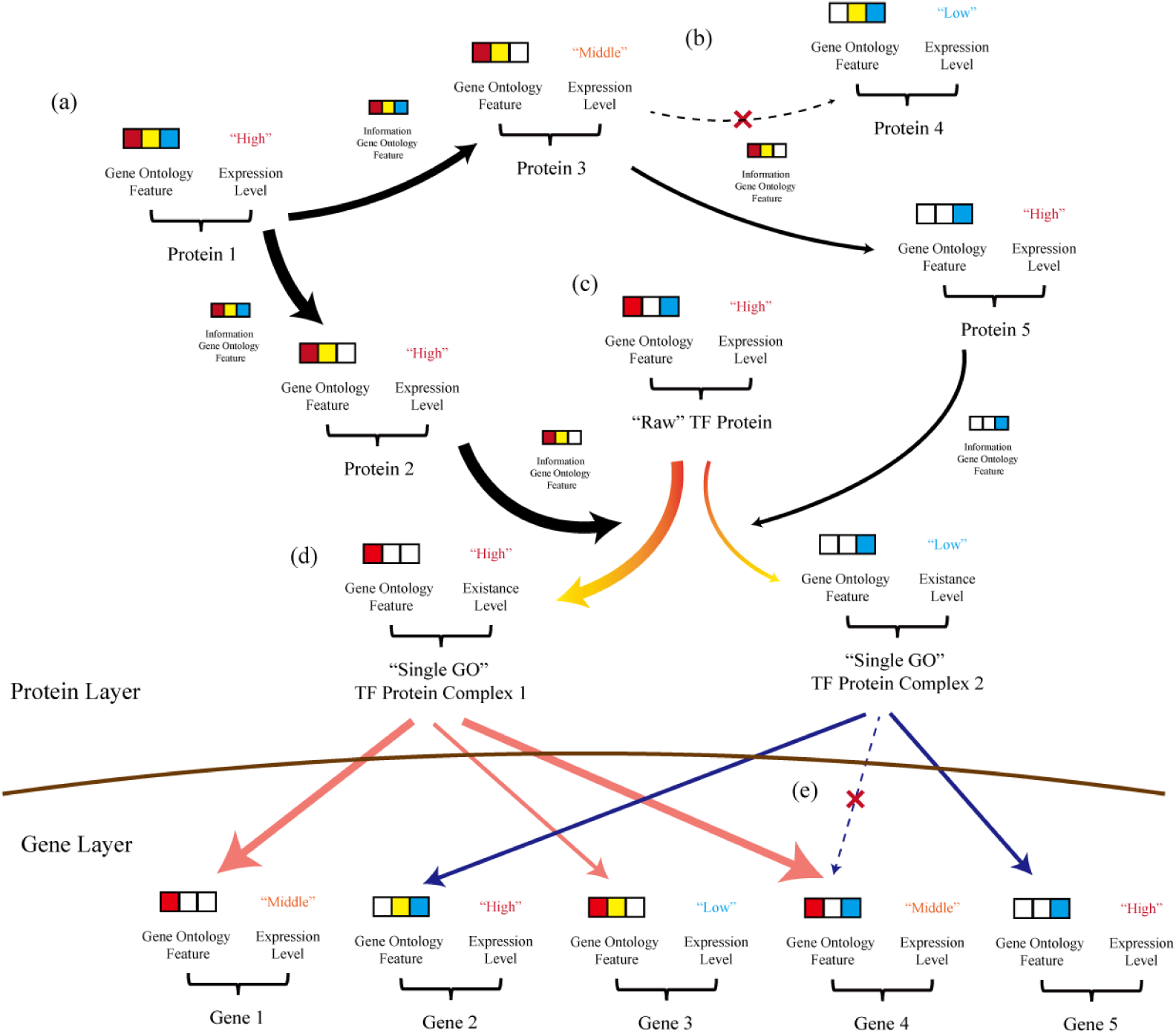
Schematic of information transmission sampling. (a) Information transmit started from “perturbation target” protein (Protein 1 here). Information was transmitted to successor proteins which were labeled “interact” with the current protein, weighted by the interaction weights from the custom PPI knowledge graph. When information was transmitted to the next protein node, the GO feature of information flux was defined as that of the current protein. (b) Only when these two proteins have correlated GO terms, the information transmission was defined to have “GO continuity”. For example, in the figure, the transmission from Protein 3 to Protein 4 had “GO continuity” while transmission from Protein 3 to Protein 5 did not. If the next protein was not a TF protein, all transmissions were allowed to ignore the “GO continuity” properties. This was a consideration of competition between different GO pathways. In the figure, the information of GO pathway Protein3-Protein4 was “grabbed” by Protein5, which had different GO properties. (c) We supposed that in most cases, the TF protein bound to DNA with the help of other proteins to form a TF protein cluster (TFpc) first. We classified the TFpcs with GO class, which was formed by the interaction of former protein with “Raw” TF proteins (e.g. Protein 2 or Protein 5 with “Raw” TF Protein). (d) Information was finally transmitted to target genes through a “Single GO” TF protein cluster, weighted by regulation weights in the custom GRN knowledge graph. (e) Only the target genes containing possible GO terms (the same or successor GO in the GO tree) were considered when calculating total sum weight. This was a simulation of selective binding of TFpcs. transmitted information with amount below a given threshold was also ignored.

**Figure 2.**
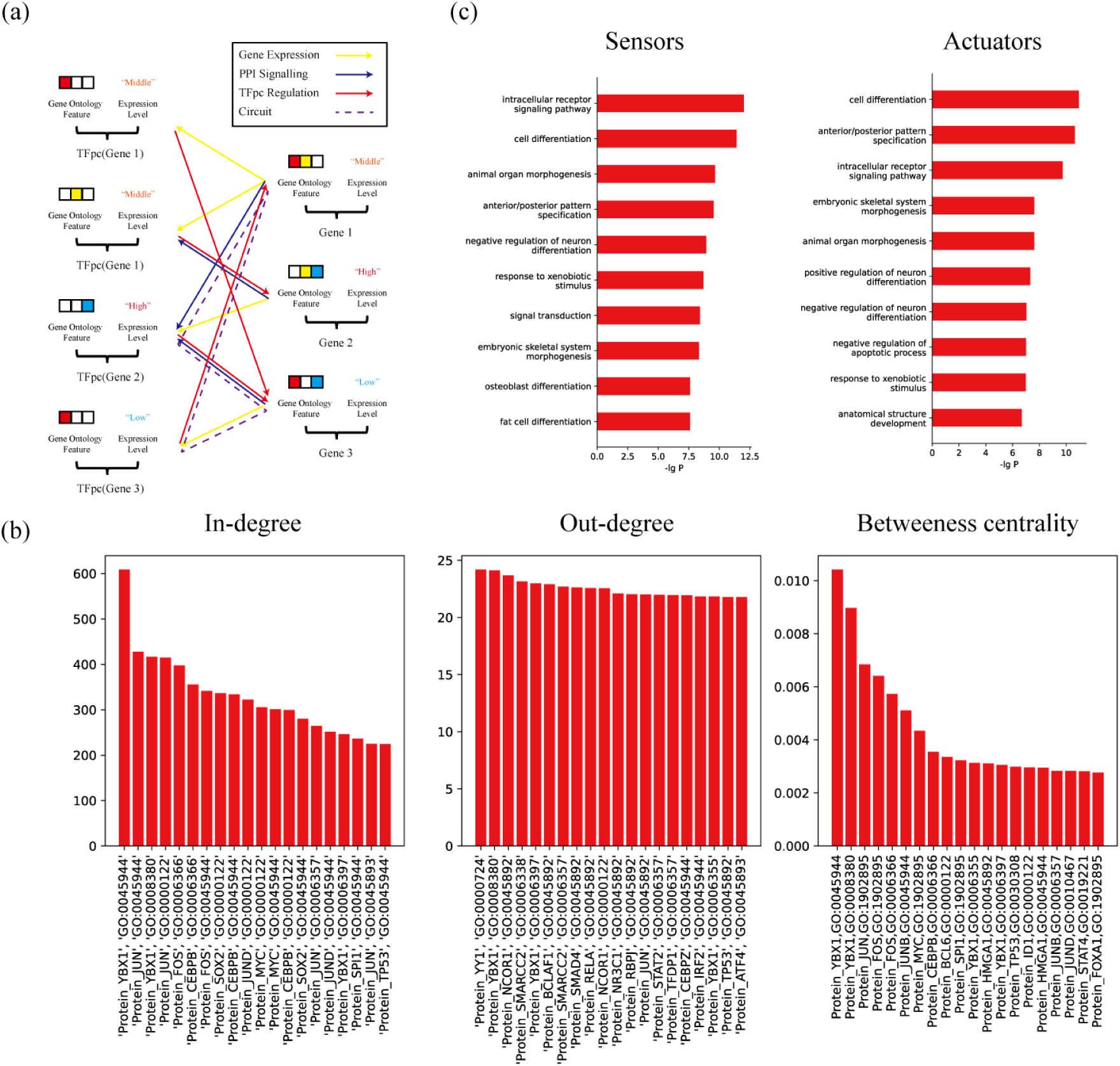
Schematic and properties of reduced regulation network. (a) Schematic of reduced regulation network. Left side: “Single GO” TFpcs. The gene that expressed the core TF protein was noted in the bracket. Right side: Genes. In the reduced regulation network drawn from information network flux (INF), there existed 3 kinds of relationships: (1) Gene expression of a TF gene to get its corresponding TFpcs (marked yellow arrows). (2) GPCI signaling from gene to TFpcs (got from INF sampling, marked blue arrows). And (3) TFpc regulation of the gene expression (got from INF sampling, marked red arrows). When there was a path from one TFpc along directed edges and finally back to the TFpc itself, we defined the path as a circuit (e.g. purple dash line in figure). (b) Physical properties of the TFpcs in the reduced regulation network (average value from all 472 cell types). Left: in-degree, Middle: out-degree, Right: betweeness centrality. (c) GO enrichment analysis of sensors (left) and actuators (right) from the union of all 472 cell types (TF-common GO terms, i.e. transcription regulation terms were ignored in this statistics).

### Properties of TFpc-Gene connection network from different cell types

We investigated the cell type-dependent networks in a tissue- and cell-type-rich database: Tabula Sapiens^30^. After constructing the networks containing both “Single GO” TFpcs and genes, we first analyzed the basic network properties. We calculated the averaged in-degree, out-degree, and betweeness centrality of “Single GO” TFpcs from each cell types (Figure 2b, SI Figure 1). On average, the reduced regulatory network contains 12,353 nodes and 98,567 edges, including 3,794 TFpc nodes and 65,569 GPCI signaling edges (SI Figure 2). The top ranked TFpcs are enriched with genes of house-keeping GO terms, which confirms their well-known biological importance. In particular, a statistics analysis after removal of the TF-common GO terms (relevant with general regulation of transcription) positions YBX1 protein cluster with GO: 0008380 (RNA splicing) on the top of in-degree, out-degree, and betweenness centrality ranking. This is consistent with its widely recognized important biological functions^31–35^.

We found that on average there are 208 sensors and 256 actuators in a particular cell type, from a total of 844 sensors and 1,183 actuators obtained for all cell types (SI Figure 3). We further performed GO enrichment for sensors and actuators in each cell type, respectively (Figure 2c). It is shown that the sensors and actuators are cell-type specific, which might be potential biomarkers for cell types. We found that in most cases, besides TF-common GO terms, both sensors and actuators enriched GO functions related to signaling and cell development, showing the importance of this feedback-circuit controlling mode in cell identity establishment.

**Figure 3.**
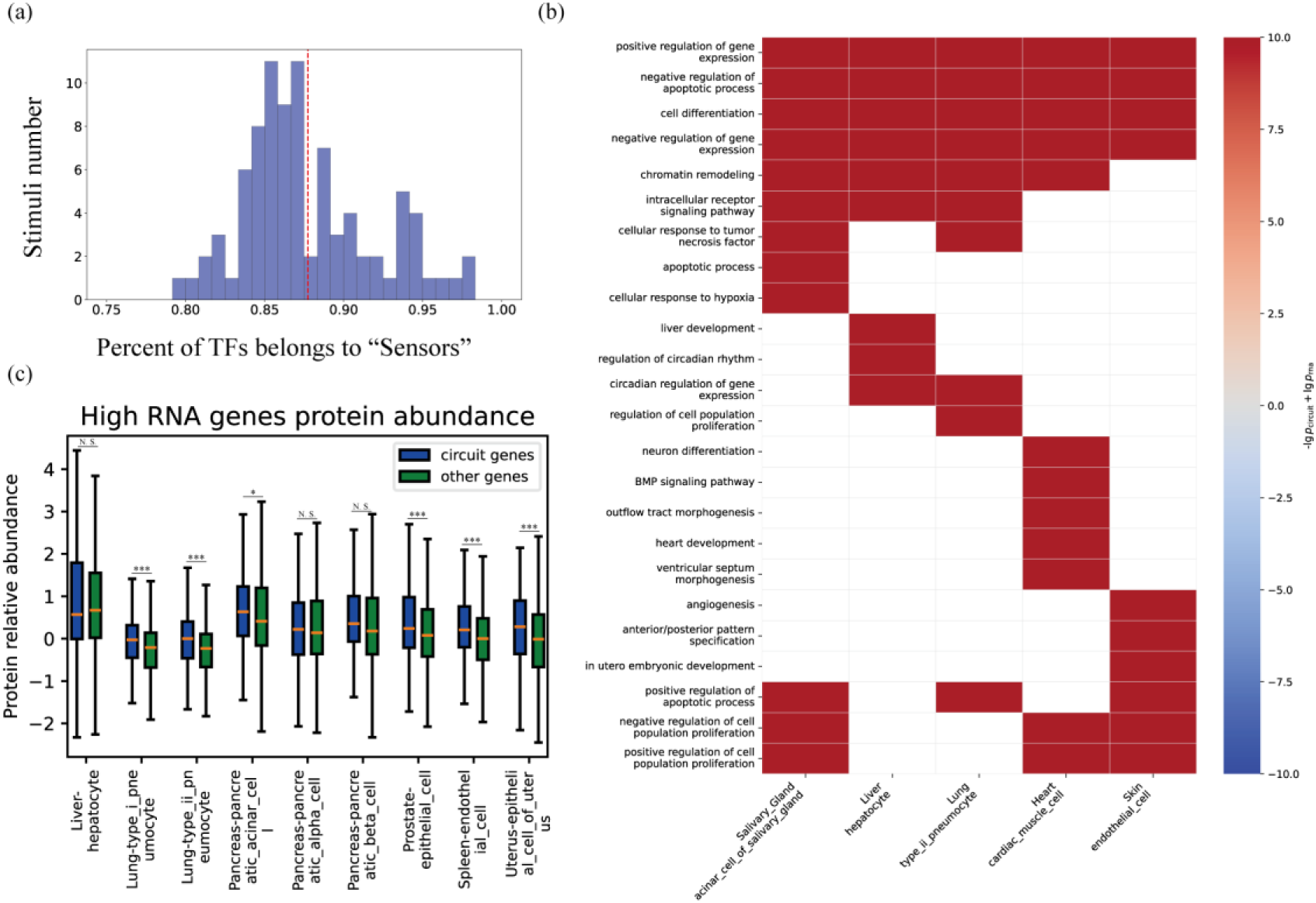
Circuits play a significant role in cellular processes. (a) Percentage of stimulation reachable TFpcs that belonged to union of sensors from 472 cell types. Results from all 89 stimuli located in the range from 0.78 to 0.99. (b) Differentially enriched GO terms between circuit genes and highly expressed genes across selected cell types. 𝑝_circuit_ and 𝑝_RNA_ are the enrichment p-values of GO terms calculated from circuit genes and from highly expressed genes, respectively. White blocks indicate that the corresponding GO terms are not enriched in circuit genes for that cell type. (c) The protein relative abundance levels of circuit genes among highly expressed genes compared with other highly expressed genes across selected cell types. Statistical significance was assessed using the one-sided Mann–Whitney U test. *** indicates p < 0.001, * indicates p < 0.05, N.S. indicates not significant.

To further validate the GO enriched function, we investigated the information transmit in response to stimuli in each cell subtype. We first combine all possible signaling fluxes to each TFpc from given perturbation source. Shortest paths to each TFpc were then identified in these information flux networks (Methods). We chose 89 stimuli from several important pathways (including WNT, TGFβ-BMP, NOTCH, Hedgehog, FGF, HIPPO, JAK-STAT, Neddylation, SUMOylation) (SI Table1). We found that the proportion of TFpcs affected by these 89 perturbation sources that belong to the ’sensor’ category ranged from 0.78 to 0.99, which indicates a “signal collection” function of these sensors (Figure 3a). This indicates the important roles of ’sensor’ TFs and their related circuits in responding to external signals.

### Function analysis of circuit genes

The existence of circuits containing more than one sensor indicates that pathways with the same or different GO terms can be coupled through the formation of circuits. To search for couplings of general biological functions, we calculated the frequency of sensor pairs that co-occur in the same circuit across 472 cell types. We found that among the most frequently occurring sensor pairs, in addition to TF-common GO terms, several GO terms associated with shared cellular functions are also coupled: For example, RFXANK and RFX5 form a circuit sharing the same GO term (GO:0045348, positive regulation of MHC class II biosynthetic process) ^36^.

We also observed that among the relatively high-frequency sensor pairs, some GO terms with similar functions are coupled: (1) The GO:0030968 (PERK-mediated unfolded protein response) pathway, regulated by ATF4, is coupled with the GO:0036500 (ATF6-mediated unfolded protein response) pathway, regulated by DDIT3^37^. (2) The GO:0006954 (inflammatory response) pathway, regulated by NFKB1, is coupled with the GO:0045087 (innate immune response) pathway, regulated by HMGB1^38^. (3) The GO:0045766 (positive regulation of angiogenesis) pathway, regulated by STAT3, is coupled with the GO:0001938 (positive regulation of endothelial cell proliferation) pathway, regulated by HIF1A^39^.

On the other hand, among the low-frequency sensor pairs, we also observed tissue-specific association of GO pathways: (1) The GO:0003253 (cardiac neural crest cell migration involved in outflow tract morphogenesis) pathway, regulated by PITX2, is coupled with the GO:0043010 (camera-type eye development) pathway, regulated by FOXC1^40^. This association specifically occurs in corneal keratocytes of the eye tissue. (2) The GO:0048289 (isotype switching to IgE isotypes) pathway, regulated by STAT6, is coupled with the GO:0006954 (inflammatory response) pathway, regulated by BCL6^41^. This latter function association specifically appears in B cells of the fat tissue.

Our findings demonstrate that the INF method not only detects established signaling pathways but also identifies correlation interactions within pathways, thereby can be used to uncover information transmission relationships in the resulting pathways that remain underexplored in current research.

### GO enrichment analysis of sensors and actuators for immune cells showed tissue-specificity

We performed a comprehensive GO enrichment analysis on immune cells across 24 tissues. Overall, the GO terms (besides TF-common GO terms) of high enrichment in these immune cells, both in sensors and actuators, are predominantly associated with signal transduction and immune-related processes, such as: GO:0140467 (integrated stress response signaling), GO:0006954 (inflammatory response), and GO:0009755 (hormone-mediated signaling pathway). We next investigated whether GO enrichment exhibits specificity for immune cells taken from different tissues. We found that GO terms with relatively low enrichment included a number of tissue-specific functions. For example, among sensors, GO:0010887 (negative regulation of cholesterol storage) is significantly enriched in the mammary gland, which is consistent with previous literature reports^42^. Another distinct example is the retinoic acid, the related GO terms (GO:0071300 (cellular response to retinoic acid) and GO:0001972 (retinoic acid binding)) of which are significantly enriched in the large intestine and prostate, aligning with their critical functions in these tissues^43,44^. Similarly, among actuators, GO:0033574 (response to testosterone) is specifically enriched in the prostate^45^, while NF-kappaB-related GO terms (GO:0032088 (negative regulation of NF-kappaB transcription factor activity), GO:0051092 (positive regulation of NF-kappaB transcription factor activity), and GO:0043124 (negative regulation of canonical NF-kappaB signal transduction)) are more broadly distributed across several tissues (prostate^46^, kidney^47^, small intestine^48^, salivary gland^49^).

### Circuit genes as potential cell type indicators

Besides sensors and actuators, other circuit genes are also expected to bear tissue-specificity. We then investigated whether the GO terms enriched in circuits across different cell types were representative of their corresponding tissue functions. For each cell type, we selected all genes included in any circuit as characteristic markers. GO enrichment analysis of these genes, compared to traits derived from highly expressed genes, revealed that circuit genes enriched more tissuespecific functions (e.g., "heart development" in cardiac muscle cells and "liver development" in hepatocytes, Figure 3b). We further validated that circuit genes indeed perform functional roles via protein expression. For each cell type, the top 10% of genes by expression level were selected and classified into circuit and non-circuit genes (Methods). By comparing their protein expression levels using bulk proteomics data from GTEx^50^ (Figure 3c), we observed that circuit genes exhibited higher protein expression in most cell types, confirming their functional importance.

The tissue-specificity of circuit genes suggests their potential utility for cell type identification. To validate this, we first examined the roles of circuit genes in cell identity establishment by using them as biomarkers for cell type identification. We further tested the utility of circuit genes as features for cell classification through a cell-typing task (Figure 4, SI Figure 4). Using the Tabula Sapiens dataset, we further clustered each cell type and selected clusters with sufficient single-cell counts as "metacells". The INF method was applied to analyze each "metacell", yielding corresponding circuit information. For comparison, we performed clustering analysis on these cell types using either all genes or only TF genes. For feature representation, we first employed a binary circuit-gene indicator (1 for circuit genes, 0 for non-circuit genes) to perform cell clustering, and compared the resulting performance with that obtained using expression-level features (Figure 4a, 4b). Both the Adjusted Rand Index (ARI) and Silhouette scores are consistently similar or higher when using the circuit-gene indicator compared to expression-level features across both gene sets (Figure 4d). We also performed metacell clustering based on expression levels using Seurat^51^ (Figure 4c). The result obtained using circuit indicators also achieved higher ARI and Silhouette scores with TFs as features, compared to Seurat (Figure 4d). Notably, clustering based on TF genes alone outperformed clustering with all genes, suggesting that sensors and actuators exhibit high cell type specificity, whereas other genes may be functionally redundant and even introduce noise into the classification task.

**Figure 4.**
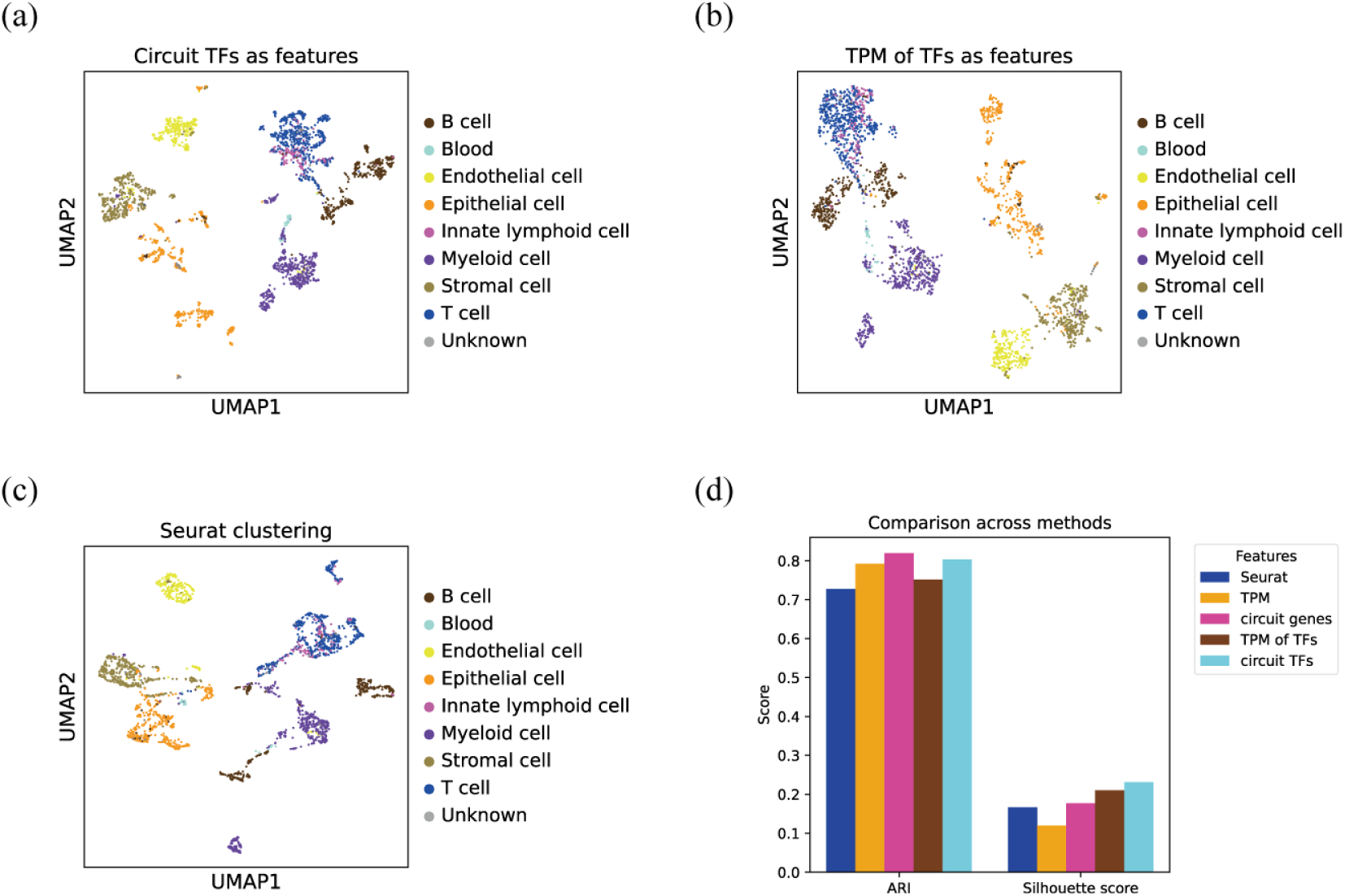
Circuit genes are better indicators for cell types. (a–c) UMAP visualization based on (a) TFs in circuits, (b) TF expression level (using transcripts per million, TPM), and (c) Seurat clustering results. Cells are colored according to major cell type clusters. (d) Quantitative evaluation of circuit genes as cell type indicators compared with expression-level-based and Seurat clustering results, assessed using Adjusted Rand Index (ARI) and Silhouette scores.

### Circuit formation and tissue-specific gene expression

Different cell types perform distinct biological functions and maintain a steady physiological state. These functions are supported by both housekeeping genes (HKGs) and tissue-specific genes (TSGs). HKGs generally exhibit stable expression across all cell types, reflecting their essential and consistent roles in basic cellular processes, whereas TSGs display high expression only within specific tissues where they execute specialized functions. To investigate how regulatory circuits contribute to these expression patterns, we analyzed the relationship between the lengths of regulatory circuits and the functional characteristics of the participating genes.

We speculate that to ensure robust and stable expression across tissues HKGs tend to participate in short but efficient regulatory circuits, whereas TSGs participate more likely in longer, tissue-specific circuits involving multiple regulatory interactions. Consistent with this hypothesis, we observed that as the number of genes within a circuit increased, the degree of tissue specificity also increased across multiple cell types (Figure 5). Moreover, the proportion of HKGs was higher in short circuits containing at most five genes, while the proportion of TSGs was higher in long circuits containing six or more genes (Figure 5). These results suggest that HKGs participate in compact regulatory circuits that facilitate efficient feedback for stability maintenance, while TSGs are regulated through extended circuits to achieve a more complex regulation and tissue-specific functionality. To illustrate this mechanism, we examined an example involving the housekeeping gene PPARA, which regulates lipid metabolism^52^. In liver hepatocytes, PPARA forms multiple regulatory circuits with NR0B2, including five long circuits that also involve BAAT, a liver-specific gene encoding an enzyme essential for bile acid metabolism^53^ (SI Figure 5). If one of these circuits is disrupted at any node other than PPARA, BAAT, or NR0B2, alternative circuits can still sustain BAAT expression, suggesting redundancy that ensures robust regulation. Additionally, NR0B2 forms a two-gene circuit with STAT3, while STAT3 forms three-gene circuits with STAT2 and JAK1, consistent with previous findings that the JAK–STAT pathway regulates lipid metabolism^54^ (SI Figure 5). Together, these results suggest that upstream HKGs form short and robust regulatory circuits with downstream transcription factors, which in turn participate in longer, tissue-specific circuits with TSGs to drive specialized cellular functions.

**Figure 5.**
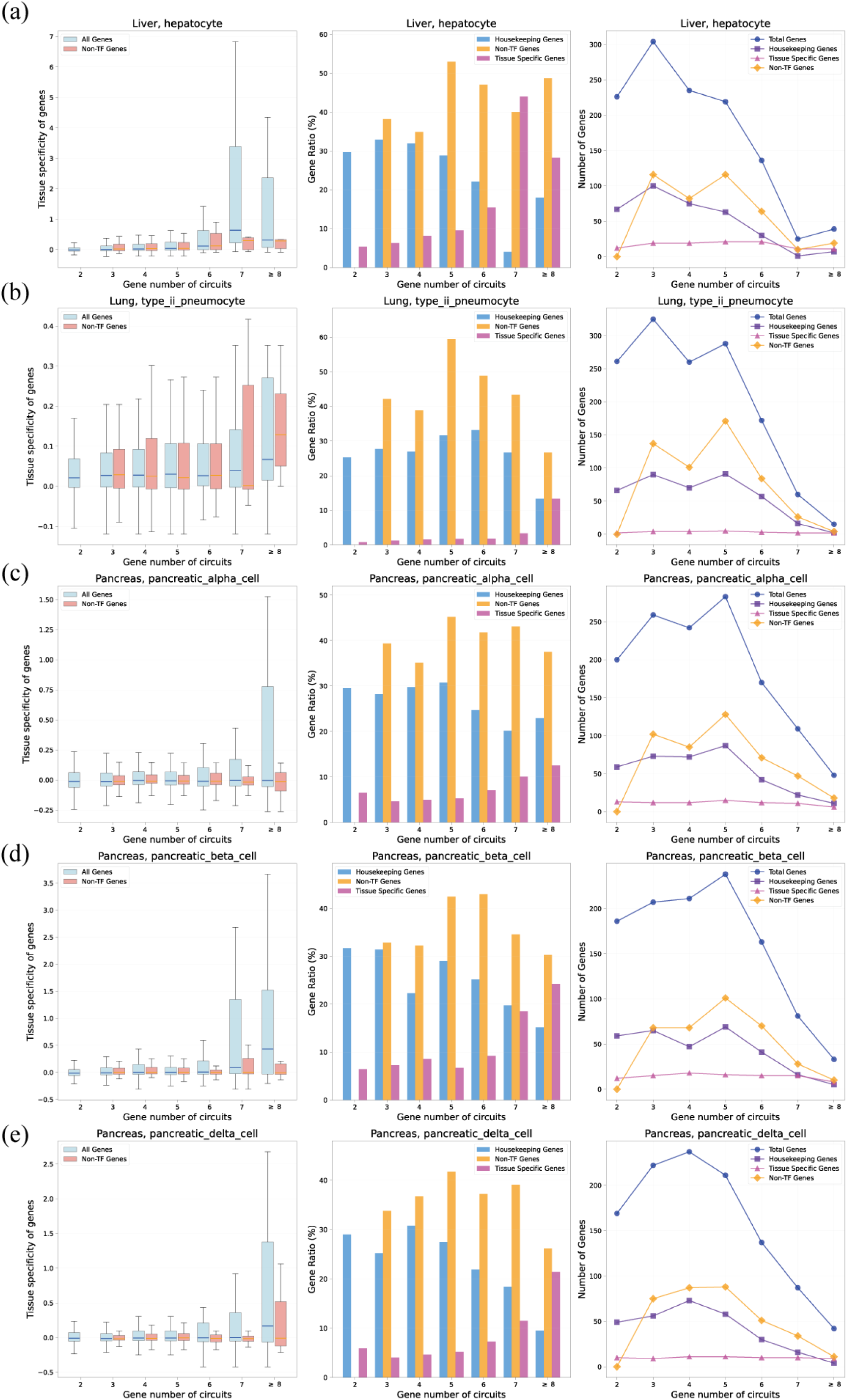
Tissue-specific genes are more enriched in long circuits. The boxplot of tissuespecificity of genes (left panel), the ratios of house-keeping genes, non-TF genes and tissuespecific genes (middle panel) and the numbers of these genes as well as total gene numbers in circuits with different gene numbers (right panel) in (a) liver hepatocytes; (b) lung type II pneumocytes; (c) pancreas pancreatic 𝛼 cells; (d) pancreas pancreatic 𝛽 cells; and (e) pancreas pancreatic 𝛿 cells.

### Investigation and theorization of the circuit-mediated signal response

The biological functions are regulated in a hierarchical way, which suggests that the circuits identified in this study should also be coupled to each other hierarchically. To systematically analyze the coupling between circuits, we first aggregated all circuits identified across the 472 tissue subtypes, resulting in a total of 276,218 distinct circuits composed of different types of genes and proteins (without distinguishing GO terms). We calculated the frequency of occurrence of each edge across all circuits. Using edges that appeared more than 10 times in total, we constructed a directed network consisting of 3,367 nodes (including genes or proteins corresponding to 858 TF genes and 1,824 non-TF genes) and 22,324 edges. Circuits that did not contain any edge from this network were defined as "periphery circuits", while those containing edges from the above network were defined as "central circuits" to emphasize the intercoupling between circuits. We identified a total of only 4,456 "periphery circuits", accounting for 1.6% of all circuits. This small portion of periphery circuits suggests that circuits are in general highly coupled, which has important implications for the robustness of cellular signaling networks.

We next proceed to investigate circuits that exclusively consist of TF gene-TF protein pairs (excluding non-TF genes). These circuits can be used to characterize the mutual regulation at the gene expression level between TFs with similar functions (both acting as sensors and actuators). We focused on the network formed by these circuits. Given the relatively small size of this network, we did not set a threshold for edges in this analysis. This network comprises a total of 1,433 nodes (including genes or proteins corresponding to 721 TF genes) and 8,833 edges. To investigate whether this network exhibits functional partitioning, we performed community analysis of this network, and obtained 17 communities. GO analysis of the 9 communities with relatively large numbers of genes showed that these communities are associated with distinct GO functions and are linked to several important signaling pathways (SI Table2). Such a segregation of regulatory circuits suggests that different functions tend to be regulated separately, to avoid severe disturbance of other functions when only one of the functional communities is perturbed. Furthermore, in individual cell types, the circuits only include a subset of edges from these communities (SI Table3), and these communities do exhibit cell type specificity (SI Table3). For example, in Community 4 (primarily associated with GO terms related to secretion and neural development), the cell types with the highest number of edges are tissues with secretion functions, such as the large and small intestines, pancreas, and eyes. As another example, the cell types with the highest number of edges in Community 6 (primarily associated with GO terms related to immunity and cell surface signaling) are predominantly immune cells in tissues closely associated with immune functions, such as blood and bone marrow. These results suggest that genes tend to organize into functional regulatory modules, and a cell’s function is highly correlated to whether, and to what extent, these modules are activated.

## [Discussion]

In this article, we developed the INF method to achieve mutation information sampling based solely on gene expression data. The fundamental framework of the INF method integrates three core networks: (1) PPI network (2) TF regulatory network, and (3) gene expression relationships. Furthermore, by integrating GO information, the INF method significantly reduces the search space at the level of TF protein regulation of downstream genes. Using AI Agents to connect users with models and tools designed for executing specific tasks is an arising trend in research^55–57^. The INF method specifically provides an explanatory tool for molecular interaction networks within virtual cell models. By enabling AI Agents to invoke the INF method within virtual cell models, knowledge acquired from large language models (LLMs) ^58–60^ and data-driven AI models can be easily integrated into the INF framework (e.g., by updating corresponding edge weights in the network or add extra PPI edges). This optimization can transform the INF into a more accurate, task-oriented explanation tool tailored for specific applications.

Through INF sampling analysis, we uncovered that circuits composed of TFpcs and their regulated genes play a critical role in extrinsic signaling transduction and cell differentiation. Current researches primarily focus on gene regulatory network (GRN), which only involves the gene layer. However, these analyses fail to explain molecular interaction mechanisms. In this study, we explicitly integrated multi-layered networks (encompassing PPI and GRN) for correlative sampling and analysis, to study the significance of protein-gene regulatory circuits and, in particular, to examine the ’sensor-actuator’ regulatory patterns within cells. Starting with highly expressed proteins as initial perturbation nodes, INF yields scores representing the informational impact from each node to TFpcs. This information passage is treated as an edge "equivalent to" those in the GRN, thereby enabling INF to proceed with its second-stage information sampling on a directed network composed of these generalized edges. This approach is taken to avoid sampling bias caused by placing individual PPI edges and more complex interactions (such as TF gene regulation involving multiple protein cooperations) on the same scale. TFpcs serve as an information transmitlayer between the GRN and PPI layers, effectively describing TFs’ signal regulatory functions in actual biological processes. In summary, the INF sampling method enhances sampling accuracy by simulating actual multi-layer signal regulation processes, laying the foundation for INF’s applications in signal transduction interpretation.

In this study, we found that cells indeed contain a large number of "sensor-actuator" regulatory pairs, which form extensive regulatory circuits. GO enrichment analysis revealed that these "sensor-actuator" regulatory pairs and circuits enrich functions of cellular signal transduction. The regulatory circuits reflect cellular states from multiple perspectives. At the gene level, circuit genes effectively reflect cell identity. In representative cell types from multiple tissues, these genes are more enriched in functions characteristic of their corresponding tissues, and those with higher transcription levels also exhibit greater protein abundance than other highly expressed genes. Moreover, the binary circuit gene indicator alone, without incorporating expression levels or network edge information, can distinguish major cell types more accurately than gene expression features. At the circuit level, HKGs and TSGs participate in circuits of different lengths: HKGs tend to participate in short, robust circuits, likely to ensure stable expression, while TSGs are more often involved in long and complex circuits, consistent with them being specifically regulated in different cell types. Above the individual circuits, they tend to assemble into functional modules, probably allowing genes and circuits of similar functions to work collectively for the robust realization of biological functions. We also found that GO pathways with different functions can be coupled through these circuits, leading to cross-talk between different functional pathways.

Apart from the aforementioned advantages, INF has certain limitations. First, the accuracy of molecular interaction network information is constrained by existing biological network knowledge, and this network is likely to be a function of cellular states. Additionally, factors such as miRNA, lncRNA, and cell metabolites should be incorporated into the INF network, although this is currently limited by restricted experimental throughput. The present protein interaction network in INF uses a directed ordinary network for description. It should be more accurately represented by a hypergraph network and represents one of the future improvement directions for INF.

In summary, we have developed the INF method, which provides a general and interpretable analytical tool for AI Agent models. This will offer significant impetus to the establishment and refinement of virtual cell models. We found circuits, which are enriched in cells and show cell-type specificity, play an important role in information transmission. Further detailed analyses on these circuits will be performed to reveal the hierarchical regulatory and organizational mechanisms that render cells their robust response to external signals while maintaining their cell identity and stability.

## [Methods]

### Preparation of pseudobulk samples

All single cells belonging to the same tissue and cell type were aggregated into a pseudobulk sample by summing the counts of each gene. The resulting count data were then transformed into transcripts per million (TPM) using the Python package RNAnorm^61^. Gene lengths were obtained from gene annotation files (.gtf) with the Python package GetGeneLength^62^.

### Construction of multi-layer network

We developed the basic gene-protein multi-layer network based on sequencing data and databases.

This network contained two main layers, which was “connected” through TFpc nodes. 18,719 gene-protein pairs were taken into analysis, which were adopted standardized names using the pyensembl^63^ (EnsemblRelease^64^ (release=100, species="homo_sapiens"))

(1) PPI layer

We combined the STRING database (only the physical interactions part)^25^ and RNA sequencing data to set the bi-directed PPI interactions. For each protein node in this network, we recorded both its normalized expression value and relative expression level (0∼10) over mean value from 24 health tissues^30^. The edges were considered to exist with probability

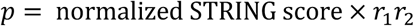

if it was in the STRING database. Here 𝑟_1_, 𝑟_2_ are relative gene expression levels of the corresponding genes. The final edge weights were set to the product of their absolute gene expression levels and above edge existence “probability” in default mode (named "p2_relascore").

(2) TF-regulation layer

For TF-regulation network, we used the gene regulatory network (GRN) skeleton from NicheNet^65^ which was read through the pyreadr^66^. For each gene node in this network, only the normalized expression value was taken into consideration. Weight of each edge was the product of the normalized expression value from nodes (default mode “both”).

In both two layers, TF nodes were marked using HumanTFDB dataset^67^.

### Construction and analysis from GO relation tree

The “go-basic.obo” file and “1GO_Gene_all.csv” file^23,24^ were downloaded from GENEONTOLOGY (https://current.geneontology.org). We only use the biological process part for analysis. To build the GO relation tree, goatools package^68^ was used, and both the relation of “contain” and “regulate” were used to set edges. When information was transmitted in INF, for each GO term in the current GO list of information flux, we concatenated it with its directed successor nodes in GO relation tree to form a “possible GO” list, and finally all these lists were concatenated together to form an “all possible GO” list. For TFpc regulation step, only when the node to be transmitted had at least one GO term in this “all possible GO” list was this transmission permitted.

### Details in Information Network Flux (INF) analysis

In most cases, we set 1,000 units of information to given perturbation node as start point. When calculating information transmitted from current node (𝐼𝑛𝑓𝑜_𝑐𝑢𝑟_) to its successor node (𝐼𝑛𝑓𝑜_𝑠𝑢𝑐_), we used the following formula:

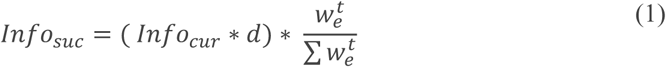

Where 𝑤_𝑒_ was the edge weight from current node to successor node, 𝑑 was the information step decay factor (default 0.85), 𝑡 was the temperature factor that controlled how sharp the probability distribution was (default 1.5). Before above transmit, if the 𝐼𝑛𝑓𝑜_𝑐𝑢𝑟_ was less than the threshold (1.0 unit of information), then this information will not transmit to other nodes and was discarded. Information was forbidden to transmit back to previous node where the information came from. When information transmitted to the TPpc node, we set half amount of it to the TPpc node (another half kept transmitting), which will be summed and used for calculation of TFpc regulation in the following step.

When calculating “Single GO” TF cluster regulation of genes, we used the following formula:

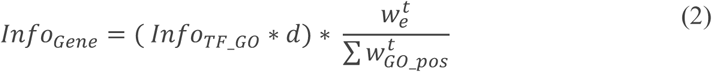

Where only when target gene’s GO terms intersect with the possible GO list that their weights were calculated.

Before sampling on DiG_signet, we first normalized its weights as follows:

1. For edges from TF genes to TFpcs with the same core TF protein and different GO terms, if they share the same sample path, we considered all GO terms to have an equal weight. The sum of all these weights from same gene was set to 0.5. This is because the same TF gene influences both the cluster of its autoregulatory TF proteins and the clusters of other TFs reached via PPI signaling. Without loss of generality, we set the proportion of its effect on the corresponding autoregulatory TF protein cluster to 0.5.
2. For edges from all expressed genes to TFpcs with a different core TF protein, the weights were normalized by the sum of weight of all edges of this kind from the corresponding genes. The sum of all these weights from the same gene was set to 0.5.
3. For edges from TFpcs to genes, the weights were normalized by the sum of weight of all edges of this kind from the corresponding TFpcs.

After sampling from all sensor and actuator candidates (similar with sampling along PPI network), we got the score of each regulation. We filtered the edges with weights below a threshold and then applied topological analysis to find circuits.

### Calculation of physical properties from reduced regulatory network

For statistics of sensor and actuator, we first analyzed from each of the 472 cell types. We set 1,000 units of information to given sensor or actuator node as start point. When calculating information transmitted from current node (𝐼𝑛𝑓𝑜_𝑐𝑢𝑟_) to its successor node (𝐼𝑛𝑓𝑜_𝑠𝑢𝑐_), we used the following formula:

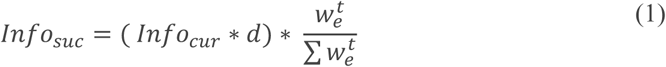

Where 𝑤_𝑒_ was the edge weight from current node to successor node, 𝑑 was the information step decay factor (default 0.85), 𝑡 was the temperature factor that controlled how sharp the probability distribution was (default 1.0). Before above transmit, if the 𝐼𝑛𝑓𝑜_𝑐𝑢𝑟_ was less than the threshold (1.0 unit of information), then this information will not transmit to other nodes and was discarded. Shortest paths and scores for possible sensor and actuator pairs were calculated. For each cell type, each cycle’s "circuit flow efficiency" score was defined and calculated as the product of score_p2g (Information transmit efficiency from sensor to actuator) and score_g2p (Information transmit efficiency from actuator to sensor), normalized by the square of initial input information amounts. The sensor and actuator list for each cell type were got from these qualified circuits. Then we summed all the sensors and actuators that were qualified in each cell type to get the total sensor and actuator box list.

The statistics of node number, TFpcs_node number, edge number, GPCI signaling edge number, TFpcs regulation edge number and gene regulation edge number were from the normed reduced regulatory network of each cell type.

### Detail methods for information transmission from stimuli

When performing INF sampling, information from given stimuli was propagated through each node and edge and finally comes to TFpcs. The total amount of information that went through each node and edge were recorded to files for each cell type. For each one of the 89 stimuli, we first calculated the averaged information amount that went through each node and edge in all 472 cell types. TFpcs with different GO were first summed to its core TF protein. Then a perturbation resource driven information flux network was generated. For each TFpc that was reachable, the shortest path from stimuli to it was found using nx.shortest_path function, with the weights of edges set to the reciprocal of total information flux.

### GO enrichment analysis of sensors and actuators of immune cells

First, we annotated the cell types according to the major clusters defined by Shi et al.^69^, with slight modifications: we added a blood cluster and merged CD4⁺ and CD8⁺ T cells into a single “T cell” cluster. The complete list of cell types and corresponding major clusters is provided in SI Table 4. T cells, B cells, innate lymphoid cells, and myeloid cells were collectively categorized as immune cells.

The efficiency of a sensor or actuator in a given cell type was defined as the maximum circuit flow efficiency among all circuits in which the corresponding protein (for sensors) or gene (for actuators) participated. Only sensors and actuators with efficiency ≥ 0.001 were retained for analysis. For each tissue, these sensors and actuators were then aggregated across all immune cell types. Then we performed GO enrichment analysis using the Python package goatools^68^. Functions with an adjusted p-value < 0.05 were considered significantly enriched. The TF list was set as the background gene list.

### Details for protein abundance analysis

We utilized the GTEx proteomic dataset^50^ to compare protein abundance between highly expressed circuit genes and other highly expressed genes. We selected the tissues that are shared in Tabula Sapiens^30^ and GTEx^50^, and chose the cell types that predominantly reflect the primary function of each tissue in Tabula Sapiens. Then we chose the genes with top 10% TPM, classified them into circuit genes and other genes, and compared the relative abundance of proteins expressed by these genes, respectively.

### Circuit genes and tissue-specific functions

To assess whether circuit genes better capture tissue-specific functions than highly expressed genes, we performed GO enrichment analysis separately for both gene sets. The method is the same as that in the method section “GO enrichment analysis of sensors and actuators of immune cells”. Five representative tissues were selected, along with the cell types that predominantly reflect the primary function of each tissue. For each circuit-gene-enriched GO term, we calculated a relative enrichment score defined as where 𝑝_circuit_ and 𝑝_RNA_ denote the enrichment p-values obtained from circuit genes and the same number of genes with the highest TPM, respectively. GO terms not enriched in highly expressed genes were assigned 𝑝_RNA_ = 1. For each of the five cell types, the top 20 enriched terms were retained. Because TFs are enriched among circuit genes, GO terms related to transcription were excluded from further analysis.

### Validation of circuit genes as potential cell type indicators

(1) Cell type clustering

To validate that the circuit genes capture cell identifications, we performed cell type clustering. We used Tabula Sapiens^30^, a single-cell RNA-seq (scRNA-seq) dataset, and compared the result obtained using circuit genes, using circuit TFs and RNA-seq data.

(2) Metacells

To reduce noise arising from the sparsity of scRNA-seq data, we aggregated single cells from the same tissue and cell type with similar gene expression profiles into a “metacell.” The circuit genes were inferred using the expression data of metacells.

To ensure the quality of metacells, we selected the cell types with at least 100 single cells. We filtered the cells with less than 100 genes and filtered the genes that were expressed in less than 3 cells. We performed single-cell clustering for the cells from the same tissue and cell type. We used the python package SCANPY^70^ for this aim:

1. Normalized the total count of different cells, with function scanpy.pp.normalize_total with default parameters.
2. Performed scanpy.pp.log1p on the scRNA-seq data with default parameters.
3. Used scanpy.pp.highly_variable_genes to choose 2000 high variable genes.
4. Performed PCA, calculated neighbors with default parameters.
5. Used Leiden algorithm^71^ with flavor = ‘igraph’, n_iterations = 2 to cluster the cells.

After cell clustering, we merged the cells in the same cluster to get metacells and calculated TPM) for all of the metacells with the same procedure as before.

(3) Metacell clustering using circuit genes

We clustered the metacells using circuit genes and circuit TFs, respectively. We used a binary circuit indicator for feature representation. If a gene (or TF) was in any circuit of this metacell, then the corresponding value of this gene (or TF) in ths metacell was 1, and 0 otherwise. The TF list was from the HumanTFDB dataset^67^. The clustering method is similar with that used when constructing metacells:

1. Filtered the genes that are circuit genes in less than 3 cells.
2. Normalized the total count of different metacells, with scanpy.pp.normalize_total with target_sum = 1e4.
3. Performed scanpy.pp.log1p on the scRNA-seq data with default parameters.
4. Performed PCA, calculated neighbors with default parameters.
5. Used Leiden algorithm^71^ with resolution = 1.0 to cluster the cells.

Then we manually annotated the metacells. To ensure that each cell type has a sufficient large number of metacells, we used major cluster labels described in the method section “GO enrichment analysis of sensors and actuators of immune cells”. A metacell cluster was assigned to a cell type major cluster if the largest number of metacells were in this cell type major cluster. If multiple major clusters had the equally largest number of metacells, then the cluster was assigned randomly to one of these cell type major clusters. The clustering results were visualized using uniform manifold approximation and projection (UMAP)^72^.

(4) Metrics

To quantitively assess whether circuit genes can infer cell identity, we used two metrics: adjusted rand index (ARI) and Silhouette score. ARI can measure the similarity between the clustering results and the ground truth annotations and was calculated using scikit-learn.metrics.adjust_rand_index^73^. The Silhouette score measures how well different cell types can be distinguished. We used the 2dimensional UMAP coordinates and the ground truth major cluster labels of metacells as inputs and calculated Silhouette score using scikit-learn.metrics.silhouette_score^73^.

(5) Baseline methods

In additional to circuit genes and circuit TFs, we also used TPM of all genes, TPM of all TFs as features, and performed the same procedure except normalization of total count. Then we compared the clustering results of different methods using the metrics.

Also, to compare with another baseline method, we performed metacell clustering using R package Seurat^51^ with TPM of all genes as features. The method is similar to that using SCANPY:

1. Normalized with SCTransform with vars.to.regress = c("nCount_RNA")
2. Performed PCA to get the first 50 PCs, calculated neighbors from these PCs with k.param = 15.
3. Performed FindClusters (using Louvain algorithm^74^) with resolution = 1.0 to cluster the cells. The annotation method of clusters method was the same as that of SCANPY. The UMAP embedding was calculated using RunUMAP with the first 50 PCs.

### Gene numbers of circuits and tissue-specificity

To investigate the relationship between circuit length and tissue specificity, we first clarified the relevant definitions. The length of a circuit was defined by the number of distinct genes it contains, with each gene and its corresponding protein counted only once. The definition of tissue-specificity is the same as that in Tian *et al.*, 2020

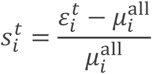

where _𝑖_^𝑡^ is the mean expression level of gene 𝑖 in tissue 𝑡, 𝜇_𝑖_^all^ is the mean expression level of gene 𝑖 across all tissues. The normalized and comparable expression data is downloaded from. Gene 𝑖 is defined as a tissue-specific gene for tissue 𝑡 if 𝑠_𝑖_^𝑡^ ≥ 2 . The house-keeping gene list are downloaded from the UCSC genome browser https://www.genome.ucsc.edu/.

### Details for community analysis

Community analysis was applied using greedy_modularity_communities function in networkx.algorithms.community, with parameters set to defaults. The version of the networkx package used in this part of the analysis was 2.8.4.

## [Acknowledgements]

This work was supported by National Natural Science Foundation of China (grant nos. T2495221, 92353304 to Y.Q.G.), Changping Laboratory (2021D-01-01) and New Cornerstone Science Foundation (grant no. NCI202305 to Y.Q.G.). We thank L. Yang, Y. Hao, W. Ma and L. Wang for technical support on computational resources. We thank Z. Ma, H. Chai, J. Wei, Z. Chen, X. Lin, L. Kong and S. Feng for helpful discussions.

## [Author contributions]

Y.Q.G. conceived the study and supervised the project. S.L. co-supervised the INF methods part. B.X. developed the INF method and implemented it. R.J. developed cell type identification method. B.X., R.J., Y.X., H.Z. performed data analysis. B.X. and R.J. wrote the initial draft of the paper. All authors contributed ideas to the work.

## [SI Figures]

**Figure S1.**
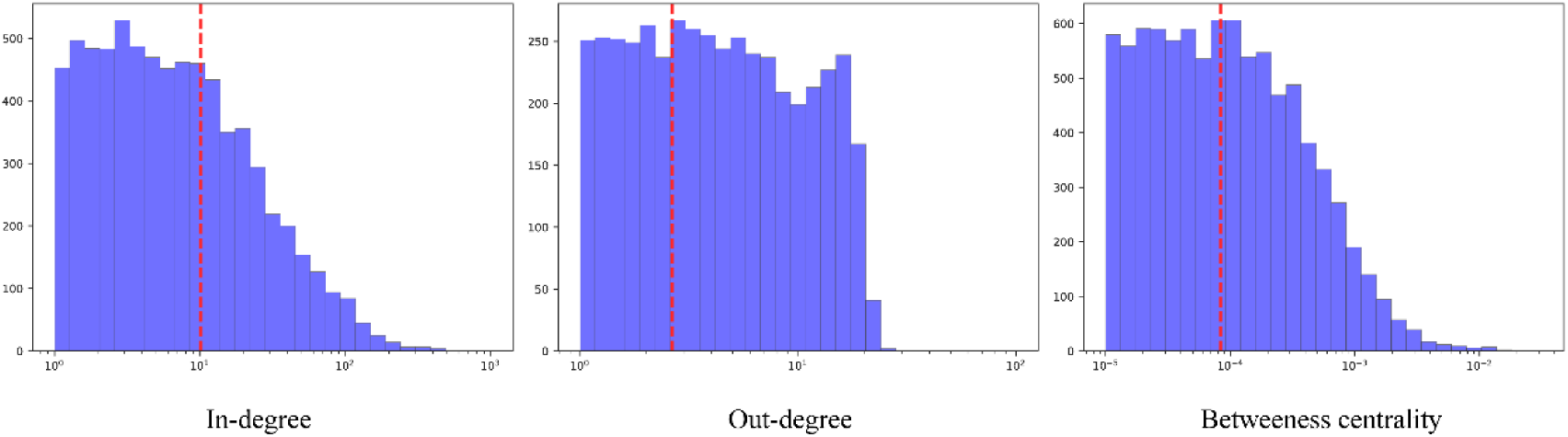
Statistics of the TFpcs’ properties in the reduced regulation network. Left: Indegree, Middle: Out-degree, Right: Betweeness centrality. Red dashed lines show the mean values: In-degree: 10.11, Out-degree: 2.65, Betweeness centrality: 8.35e-5. Through statistics on 472 tissue subtypes, we observed that TFpcs have a total in-degree of 111,813 and an outdegree of 29,281, indicating that compared to transmitting signals to downstream target genes, TFpcs possess more channels for receiving external information from the protein level

**Figure S2.**
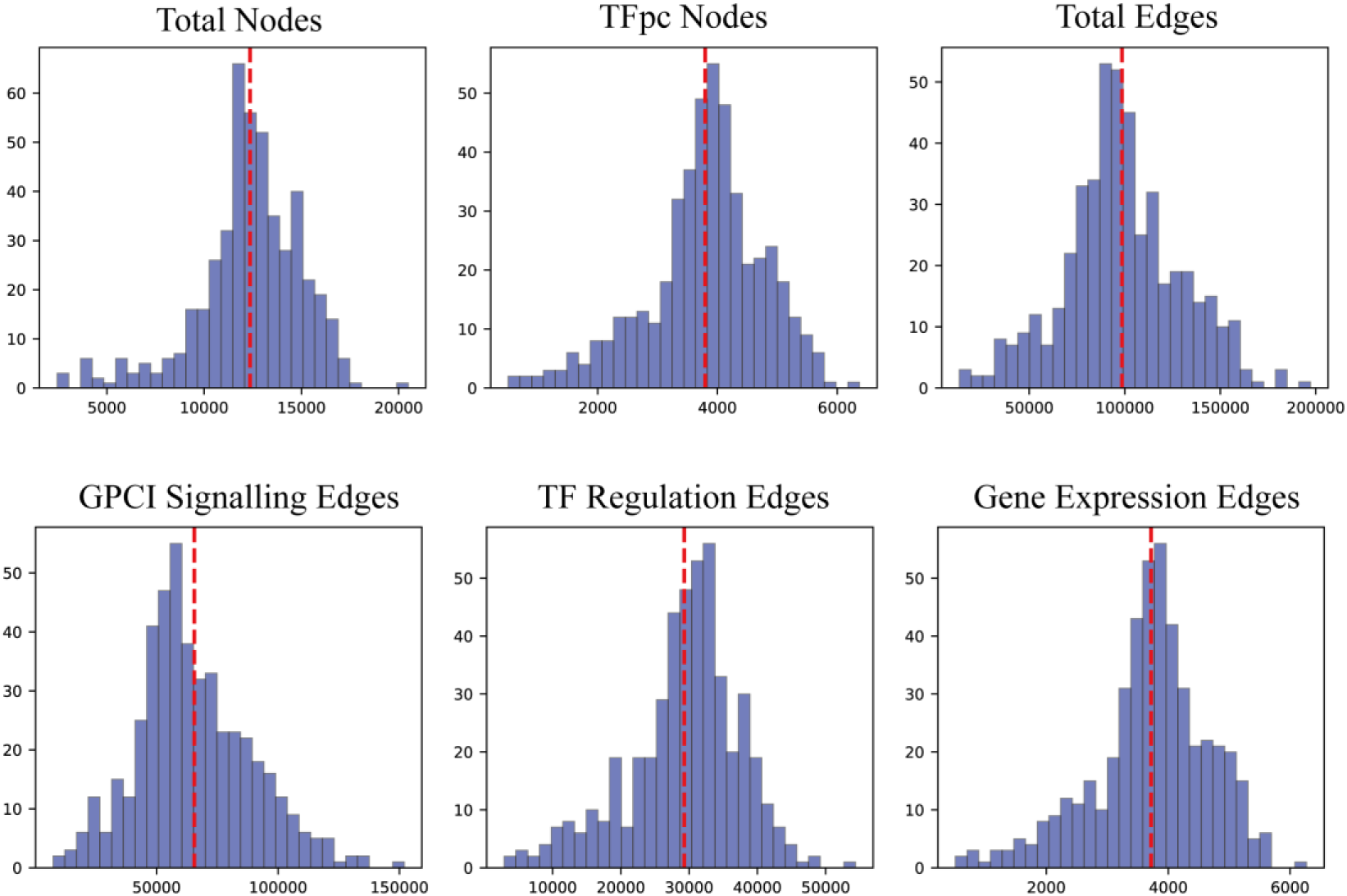
Distribution of node and edge numbers in each cell type. Histogram shows the cell type counts distribution of total nodes, TFpc nodes, total edges, GPCI signaling edges, TF regulation edges and gene expression edges (statistics on 472 cell types in total). Red dashed lines show the mean values: total nodes: 12,353, TFpc nodes: 3,794, total edges: 98,567, GPCI signaling edges: 65,569, TF regulation edges: 29,281 and gene expression edges: 3,716.

**Figure S3.**
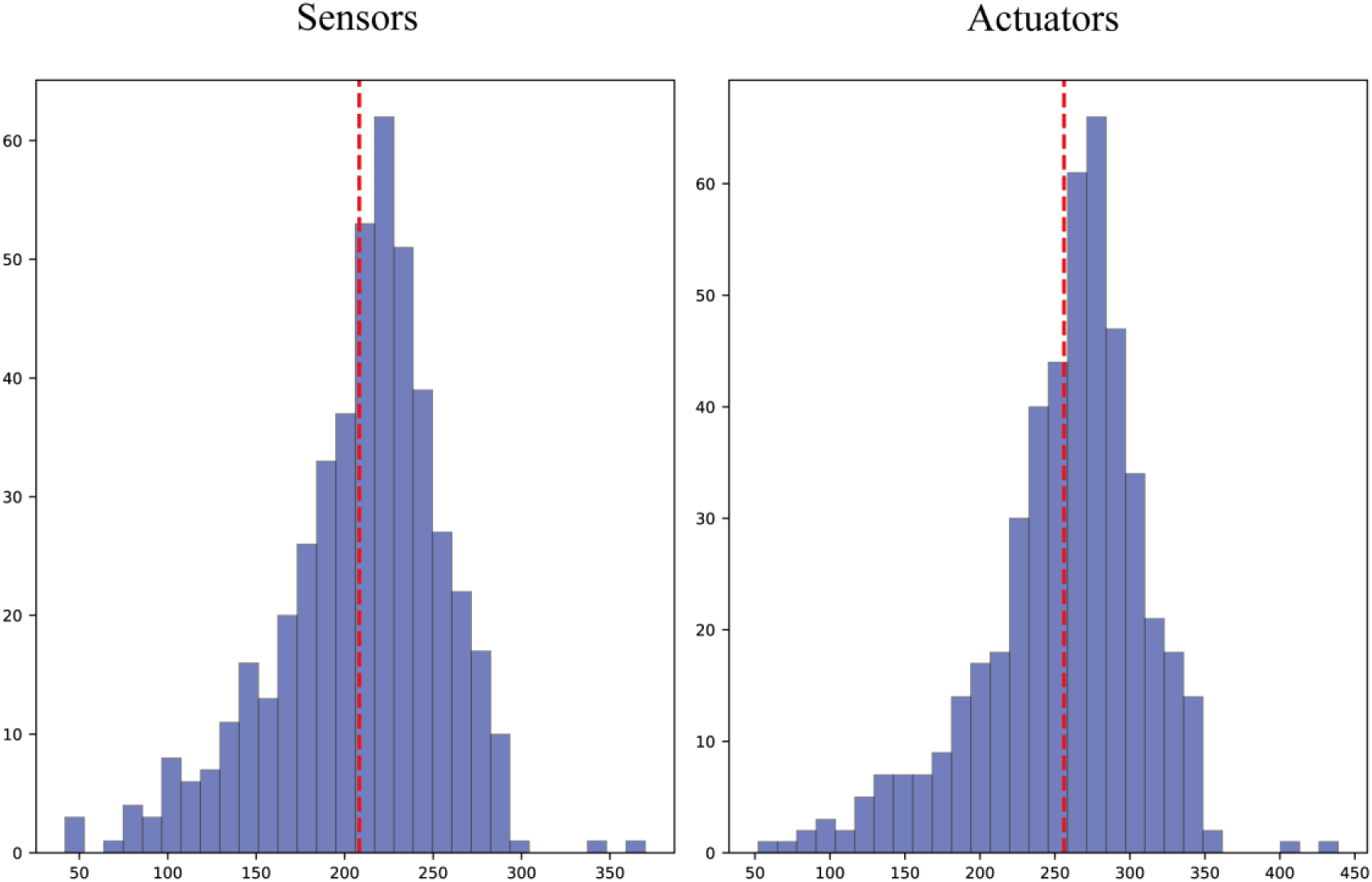
Distribution of sensor and actuator numbers in each cell type. Histogram shows the cell type counts distribution of sensor and actuator numbers (statistics on 472 cell types in total). Red dashed lines show the mean values: Sensors: 208 Actuators: 256.

**Figure S4.**
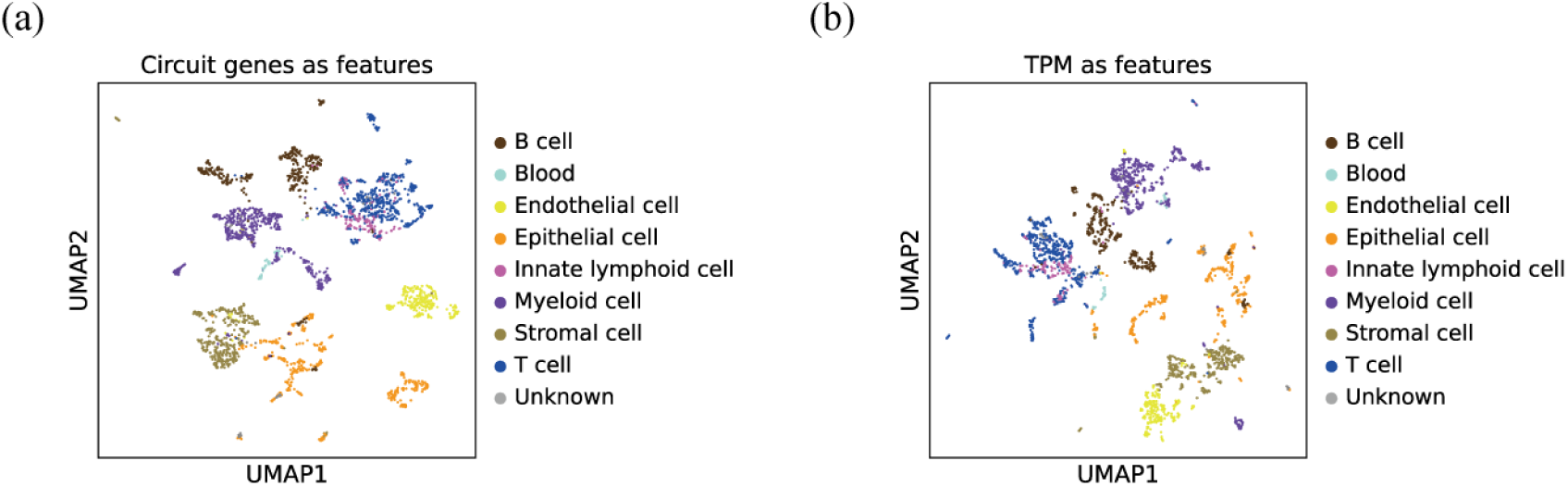
Results of alternative feature sets used for cell type identification. UMAP visualizations based on (a) circuit genes, (b) gene expression levels (transcripts per million, TPM). Metacells are colored according to major cell type clusters.

**Figure S5.**
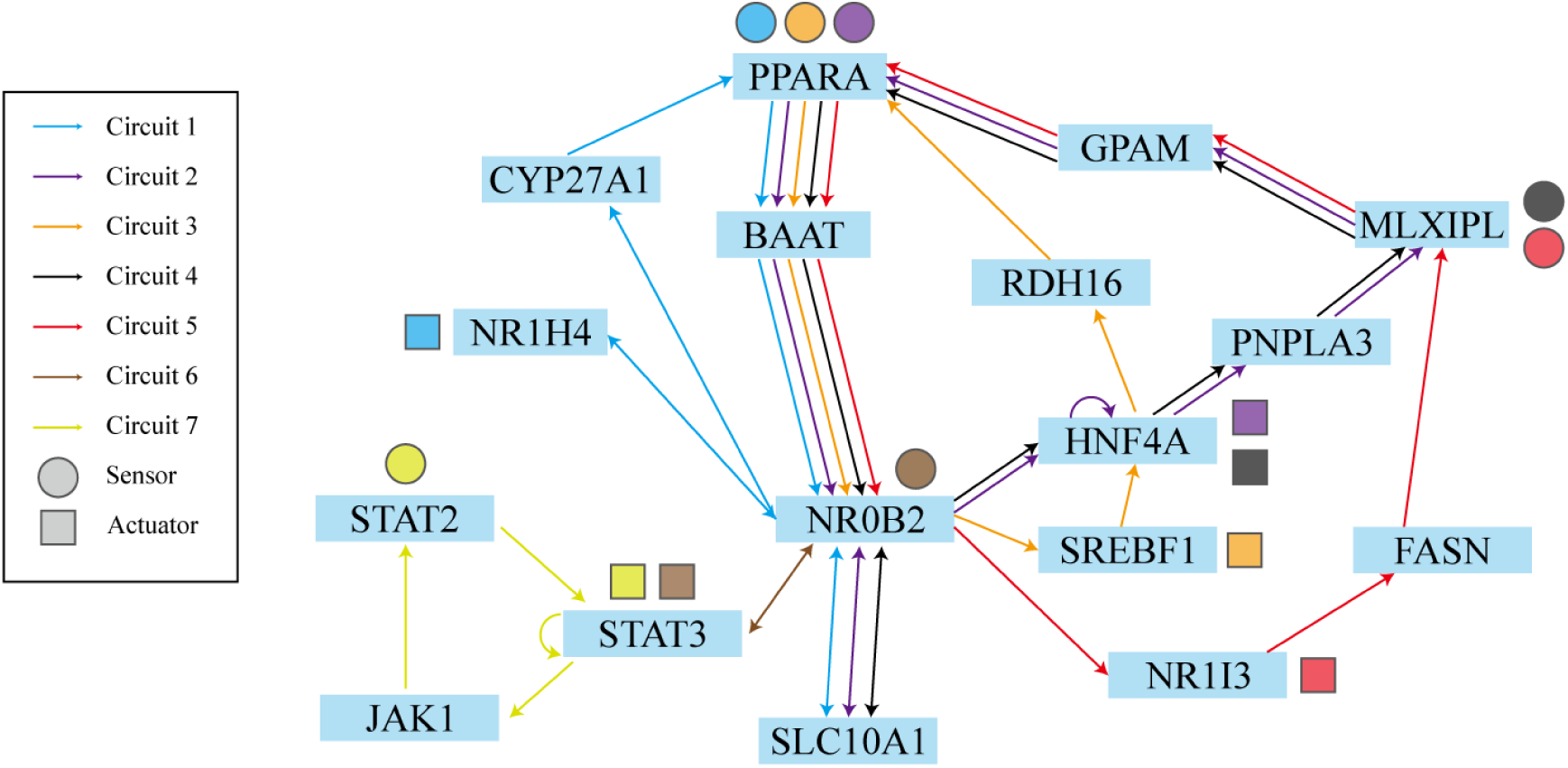
**Graphic illustration of circuits with BAAT gene (a Liver-specific gene) in liver hepatocytes**

